# Talin1 adhesions’ morphology is largely unaffected by polyacrylamide substrate stiffness

**DOI:** 10.1101/2025.04.30.651087

**Authors:** Joanna Hajduk, Patrycja Twardawa, Zenon Rajfur, Zbigniew Baster

## Abstract

Cells sense the stiffness of their extracellular matrix (ECM) and adapt their behavior accordingly. We investigated how ECM stiffness affects the spatial organization of talin1, a key mechanosensitive focal adhesion protein. Using polyacrylamide (PA) hydrogels with tunable stiffnesses (0.2–188 kPa), we analyzed cell morphology, migration, talin1 distribution, colocalization with tensin3, and fibronectin deposition.

Softer substrates enhanced filopodia activity and altered migration behavior. On softer ECMs, talin1 displayed a more even intracellular distribution, whereas on stiffer matrices it localized to the cell periphery. PA gels supported elongated talin1-based adhesions, whose morphology showed minimal variation across the 3–188 kPa stiffness range. Talin1–tensin3 colocalization was maintained regardless of stiffness, indicating a stable interaction. Notably, cells deposited more fibronectin on softer substrates.

While talin1 adhesion morphology varied little with stiffness, cell migration behavior changed markedly. Combined with prior studies, our data suggests that ECM stiffness regulates talin1 primarily through conformational changes rather than macroscopic adhesion remodeling. These findings highlight talin1’s central role in translating mechanical cues into dynamic cellular responses.

**Summary statement:** Talin1 forms elongated adhesions and robustly colocalizes with tensin3 across varying matrix stiffnesses, showing that their spatial organization is largely insensitive to mechanical cues.

## Introduction

The rigidity of the extracellular matrix (ECM) influences cell morphology and behavior. The first studies concentrating on this phenomenon were pioneered in the late 90s of the past century by the group of Yu-Li Wang (Dembo and Wang, 1999; Lo et al., 2000; Pelham and Wang, 1997). Since then, the research community has started exploring the new field of mechanobiology, aiming at uncovering molecular mechanotransduction pathways mediating cellular response to mechanical cues and looking for applications of the newly acquired knowledge to develop new, better therapies (Nelson et al., 2024).

One of the broadly studied aspects since the onset of mechanobiological research has been cell-ECM adhesion. Multiple studies have demonstrated the significance of adhesion proteins in various processes, such as embryological development (Camp et al., 2024; Monkley et al., 2011), immunological responses (Feng Lim et al., 2020; Jaumouillé and Waterman, 2020), or cancer invasion and metastasis (Baster et al., 2020; Baster et al., 2025). By taking a closer look into different cellular adhesion structures, we can distinguish several main classes: young nascent adhesions (<0.25 µm in diameter), which represent the initial stage of adhesion formation; developing focal complexes (∼0.5 µm); focal adhesions, which are mature, usually dynamic structures, involved in transducing the highest forces and facilitating migration (1 - 5 µm in length); and stable fibrillar adhesions, the most mature, centrally located structures that anchor cells to the substrate and promote ECM remodeling (>5 µm) (Gardel et al., 2010). Nascent adhesions, localized near the leading edge of cellular protrusions, undergo rapid assembly and disassembly, with an average lifetime of around 80 seconds. They can mature into focal adhesions, which are more stable structures with an average lifetime of 20 minutes. Stable fibrillar adhesions can develop from focal adhesions (Gardel et al., 2010). Molecular studies have shown that the formation, maturation, and identity of adhesion structures are tightly connected with protein composition, their posttranslational modification, and even conformation changes resulting from intracellular forces acting on proteins (Goult et al., 2018; Han et al., 2021; Vicente-Manzanares and Horwitz, 2011). One prominent example of a force-dependent adhesion-regulating protein is talin1. It serves as a crucial linker between integrins and F-actin, undergoing conformational changes in response to mechanical force. (Goult et al., 2018; Kanchanawong et al., 2010). Under tension, talin1 can stretch and reveal cryptic binding sites for vinculin, enhancing the integrin–cytoskeleton connection and facilitating focal adhesion maturation (Goult et al., 2013). Conversely, reduced forces or altered molecular interactions may modify talin1’s conformation and localization within adhesions (Han et al., 2021; Kumar et al., 2016). This mechanosensitive behavior of talin1 is thought to tune adhesion composition in response to physical cues, establishing talin1 as a mechanosensitive signaling hub (Goult et al., 2018). Moreover, talin1’s role extends beyond focal adhesions at the cell periphery, as it can also be found centrally in lower abundance (Praekelt et al., 2012).

Tensin-family adhesion proteins are primarily associated with a distinct class of adhesions – fibrillar adhesions – that form centrally as cells pull and remodel fibronectin (Katz et al., 2000). Tensin3, in particular, is predominantly localized to fibrillar adhesions (Clark et al., 2010). Recent work by Atherton and colleagues (Atherton et al., 2022) demonstrated that talin1’s interaction with tensin3 drives the formation of fibronectin-associated fibrillar adhesions. This finding suggests an unexpected crosstalk between the talin-mediated force transduction machinery and the tensin-mediated matrix assembly pathway. Indeed, fibrillar adhesions themselves have been shown to be mechanosensitive (Atherton et al., 2022; Barber-Pérez et al., 2020), yet the results seem to be contradictory, highly dependent on analysis method, thus requiring further investigation.

While several studies have examined the recruitment of other adhesion proteins (such as paxillin and tensin) in response to matrix stiffness (Barber-Pérez et al., 2020; Oakes et al., 2018; Pelham and Wang, 1997), the mechanosensitive reorganization of endogenous talin1 has not been explored in depth. Here, we focused on how ECM stiffness affects talin1 organization in mouse embryonic fibroblast cells, using polyacrylamide (PA) gels to simulate varying tissue elasticities. First, we examined how the mechanical properties of the culture substrate influence cellular morphology and migration modes. Next, we analyzed talin1 distribution in cells cultured on substrates of different stiffnesses and observed the formation of elongated talin1-based structures at the ventral membrane regardless of substrate stiffness. To further contextualize our findings, we analyzed the correlation between talin1 and tensin3. Interestingly, while this correlation was independent of ECM stiffness, we noted distinct differences between glass and polyacrylamide substrates. Finally, we assessed fibronectin secretion and found an inverse relationship with substrate stiffness.

## Results

### ECM stiffness influences mouse embryonic fibroblasts’ morphology and migration pattern

Pelham and colleagues (Pelham and Wang, 1997) and later Oakes and colleagues (Oakes et al., 2018) showed that in MEF cells, soft ECM induces cell edge fluctuations and changes their migration pattern from the most common for these cells, the lamellipodial mode, to the filopodial mode. In our initial step, we analyzed the dependence of cell morphology and migration mode of NIH/3T3 Tet-Off stably expressing mCherry (not regulated by TRE promoter) cells on ECM stiffness. We plated cells on PA substrates of a range of 0.2 kPa – 195 kPa of stiffnesses coated with fibronectin (10 µg/ml) and let them adhere for 24 h. We also included cells seeded on fibronectin-coated glass as a control. We imaged cells using a high NA lens (1.46 NA, Apochromat) in DIC contrast, which allowed us to observe fine morphological and migratory details. We took up to 30 min time-lapse movies with 10 s intervals (Movies 1-10). Cells plated on glass and 18 - 188 kPa substrates showed similar morphology and migration patterns (Movies 1-5). Cells on intermediate stiffnesses, 0.5 – 11 kPa, had reduced spreading, produced multiple filopodia or filopodia-like protrusions, and an altered migration pattern resembling filopodial or filopodia-lamellipodial mode (Movies 6-9). These cells resembled those plated on 2.1 kPa polyacrylamide substrates in the work of Oakes and colleagues (Oakes et al., 2018). Cells on the softest, 0.2 kPa, substrates were rounded up and usually unable to migrate. They also produced a limited number of filopodia (Movie 10).

### Talin1 distribution depends on ECM stiffness

In their studies, Oakes and colleagues (Oakes et al., 2018) compared paxillin distribution between NIH/3T3 cells plated on polyacrylamide substrates of 2.1 kPa and 48 kPa. Similarly, Barber-Pérez and colleagues (Barber-Pérez et al., 2020) studied the distribution of fibrillar adhesion-related protein tensin1 in cells cultured on substrates with a stiffness gradient. There are a couple of other studies showing that the recruitment and distribution of adhesion proteins in cells are dependent on the mechanical properties of the ECM (Guo et al., 2006; Han et al., 2021; Pelham and Wang, 1997; Zhou et al., 2017), but, to our knowledge, none of them studied the distribution of endogenous talin1 protein in-depth.

In our studies, we used the 97H6 mouse anti-talin1 monoclonal antibody. We specifically opted for this clone because, to our knowledge, at the time of the study, it was the only commercially available talin1-specific antibody detecting the mouse protein variant. As previously, we plated NIH/3T3 cells (without Tet-off and mCherry inserts) on fibronectin-coated (10 µg/ml) glass and a selected range of polyacrylamide substrates of varying stiffnesses (3 kPa, 18 kPa, 29 kPa, and 188 kPa) for 20 h. Then, we fixed them in freezer-cold methanol (−20°C), stained for talin1, and imaged in 3D using a laser scanning confocal microscope.

First, we analyzed the distribution of talin1 at the ventral membrane of the cell (Fig. 1,2). We divided cells into 1 µm regions depending on the distance to the edge (Fig. 1B) and quantified the average intensity for each area (Fig. 1A-B). For further analysis, we included data only from the first 10 µm from the edge or 2/3 of the total distance from the edge, whichever was smaller. We normalized data from each cell to the highest value, averaged them, and plotted normalized intensity vs distance from the edge for each condition (Fig. 2A). We noticed that the distributions resembled exponential decay curve; thus, for quantitative measurements, we fit exponential curves using all data points for each condition and compared fitted models for the fit of plateau levels and exponential coefficients (Fig. 2B-C).

**Fig. 1.**
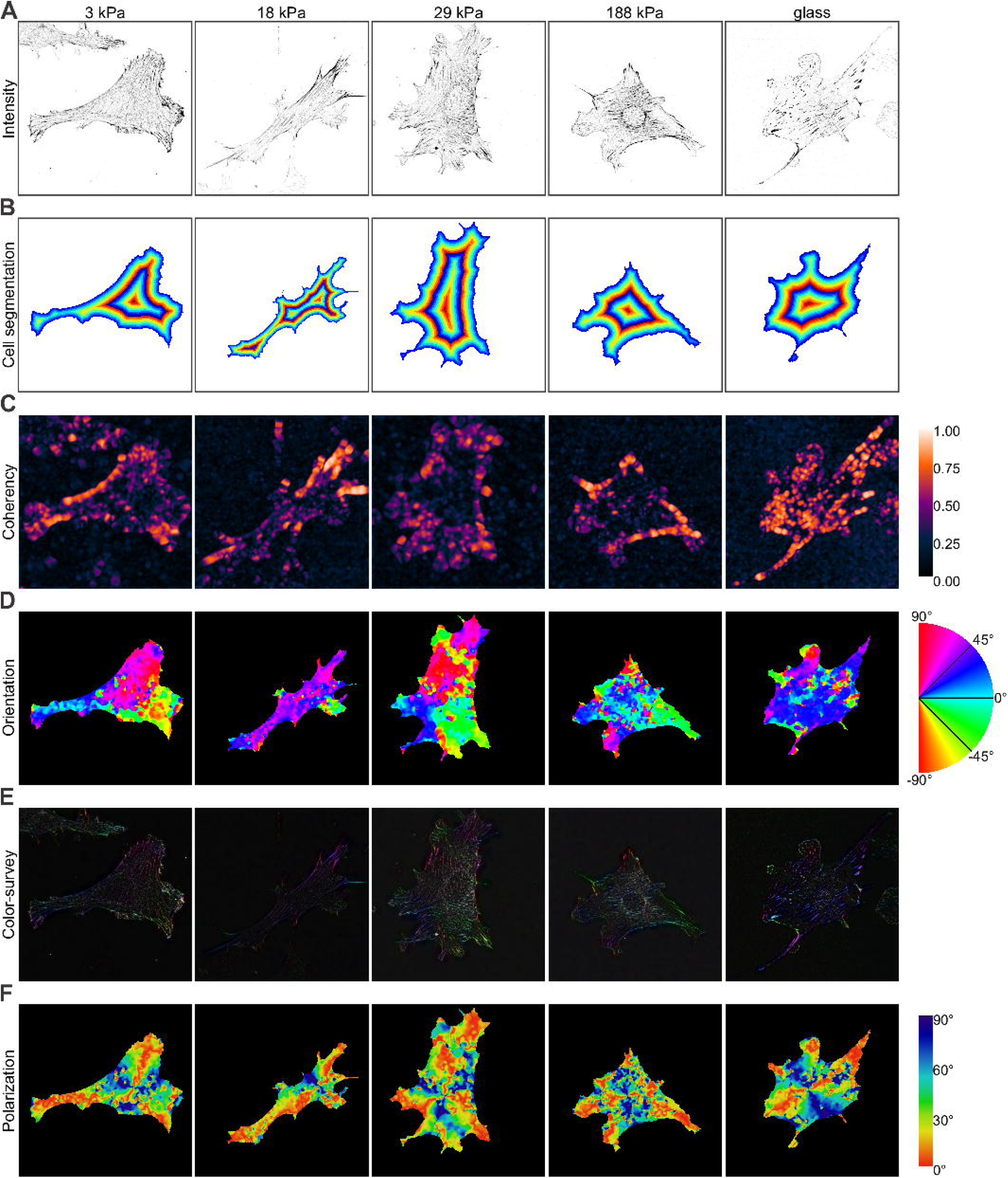
Talin1 distribution, coherency, and orientation depending on ECM stiffness. **A**. Representative images of spatial distribution of talin1 in cells plated on glass or elastic PA gels of different elasticity (3-188 kPa) (scale bar: 20 µm). Single planes focused at ventral membrane were selected from stacks, then noise was removed with N2V algorithm, and background was removed using rolling ball method (paraboloid, rolling radius 1 px). **B**. Cell segmentation into 1 µm thick outlines. Each color represents a different outline. **C**. Coherency of talin1 distribution, with 0.5 µm window of analysis. **D**. Color-coded orientation of talin1 adhesions, with 0.5 µm window of analysis. **E**. HSB mode superposition of talin1 intensity (brightness), coherency (saturation), and orientation (color, as in D). **F**. Deviation from radial polarization of talin1 adhesions, based on D (0° - perfectly aligned).

**Fig. 2.**
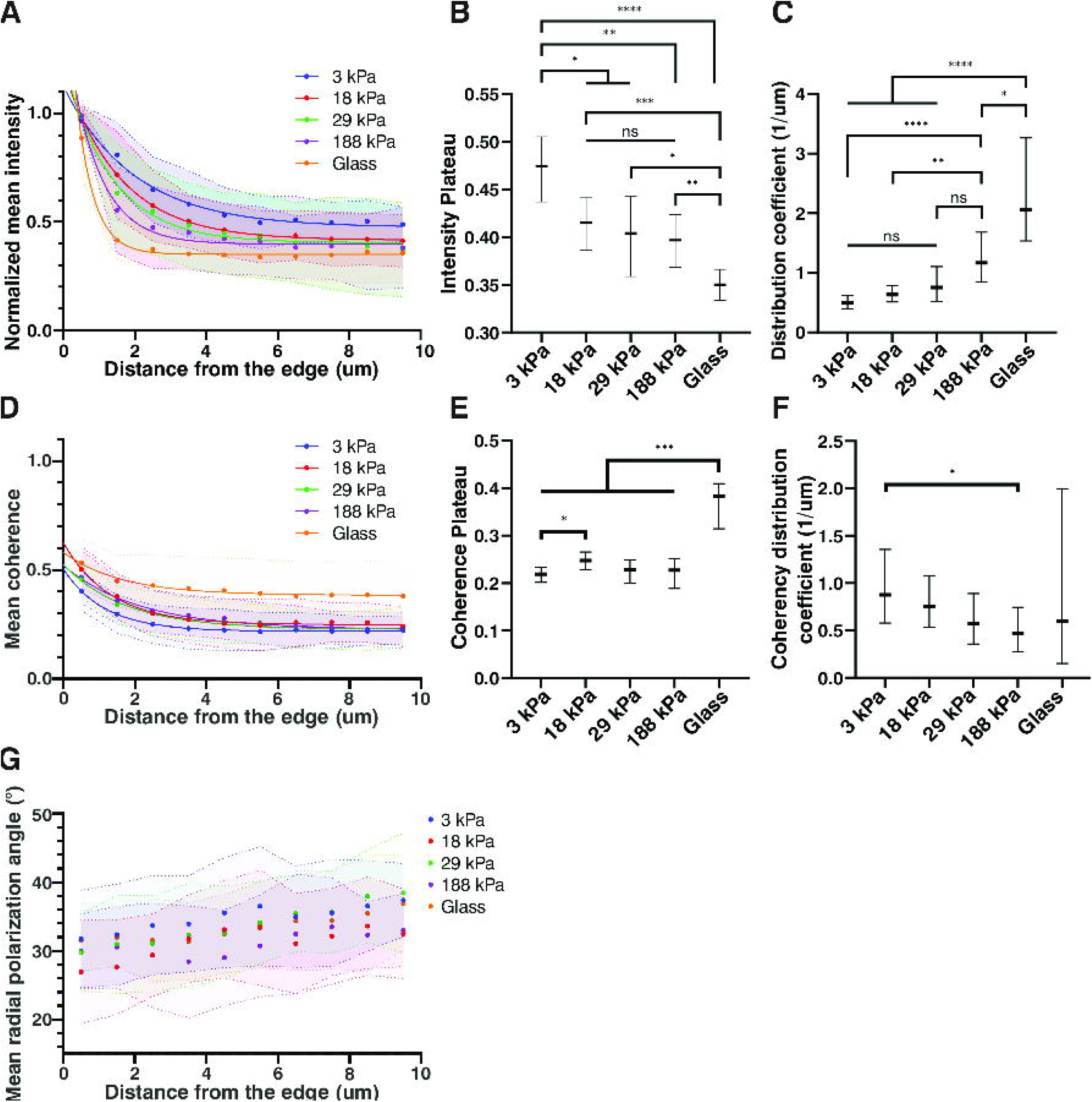
Quantification of talin1 distribution, coherency, and orientation depending on ECM stiffness. **A**. Distribution of talin1 in cells based on distance from the edge, with exponential fitting to the data. Based on Fig. 1A-B. **B**. Comparison of intensity plateaus of the fits to talin1 distributions in A. **C**. Comparison of exponential coefficients of the fits to talin1 distributions in A. **D**. Distribution of talin1 distribution coherency in cells based on distance from the edge, with exponential fitting to the data. Based on Fig. 1B-C. **E**. Comparison of intensity plateaus of the fits to talin1 distribution coherencies in D. **F**. Comparison of exponential coefficients of the fits to talin1 distribution coherencies in D. **G**. Distribution of talin1 radial polarization in cells based on distance from the edge. Based on Fig. 1B,F. Number of repeated experiments: 3. Total number of cells analyzed: 3 kPa – 33, 18 kPa – 31, 29 kPa – 26, 188 kPa – 25, glass – 28. Statistical analysis: Extra sum-of-squares F test for single parameter comparison of fits. * p ≤ 0.05, ** p ≤ 0.01, *** p ≤ 0.001, **** p ≤ 0.0001.

We observed higher plateau levels for softer ECM levels than for stiffer ones (Fig. 2B). It is important to note that due to the normalization, we cannot definitely state talin1 levels are higher in the center in cells cultured on softer substrates, but rather, talin1 is more evenly distributed throughout the cell. The comparison of exponential coefficients showed higher values for stiffer substrates, indicating a steeper decrease in talin1 concentration in these substrates (Fig. 2C).

Next, we analyzed coherency and radial polarization towards the center of the cell of talin1 using OrientationJ Fiji plugin (Püspöki et al., 2016; Schindelin et al., 2012) (Fig. 1C-F). In the plugin, we applied 0.5 µm window for analysis. We divided and processed the data in a similar way to how we did with talin1 distribution, using intensity as weight for averaging. Coherency at plateau for glass was significantly higher than for elastic substrates, but we see only a little difference between PA substrates (Fig. 2E). Coherency showed slightly higher values closer to the cell edge than to the center for all conditions (Fig. 2D). On PA substrates exponential coefficient increases with decreasing stiffness, yet our data in most cases does not show significance (Fig. 2F). Glass and all PA substrates showed average radial polarization of talin1 at a level of 30°, diverging towards random organization at the center (0° - aligned with radials; 45° - random or diagonal organization; 90° - perpendicular to radials). We did not see any correlation with substrate type or stiffness (Fig. 2G).

### Talin1 adhesions’ morphology exhibits minimal variation across polyacrylamide substrate stiffnesses

Next, we analyzed the morphology of talin1-associated adhesions in relation to ECM stiffness using the same images. For this purpose, we developed adhesion analysis software (Twardawa et al., *in prep*). We initially expected to find smaller, rounded-up talin1-associated adhesion structures in cells plated on soft substrates, similar to those described for other adhesion proteins shown in other works (Han et al., 2021; Oakes et al., 2018; Pelham and Wang, 1997; Plotnikov et al., 2012). Surprisingly, on PA gels, we mainly found slimmer, elongated structures, often ranging throughout the whole cell length with only slight responsiveness to PA stiffness (Fig. 3A). We compared the structures quantitatively using, among others, area, axial ratio, and circularity parameters. Axial ratio refers to the proportion between the longest and shortest axes of a structure. Circularity measures how closely the structure resembles a perfect circle.

**Fig. 3.**
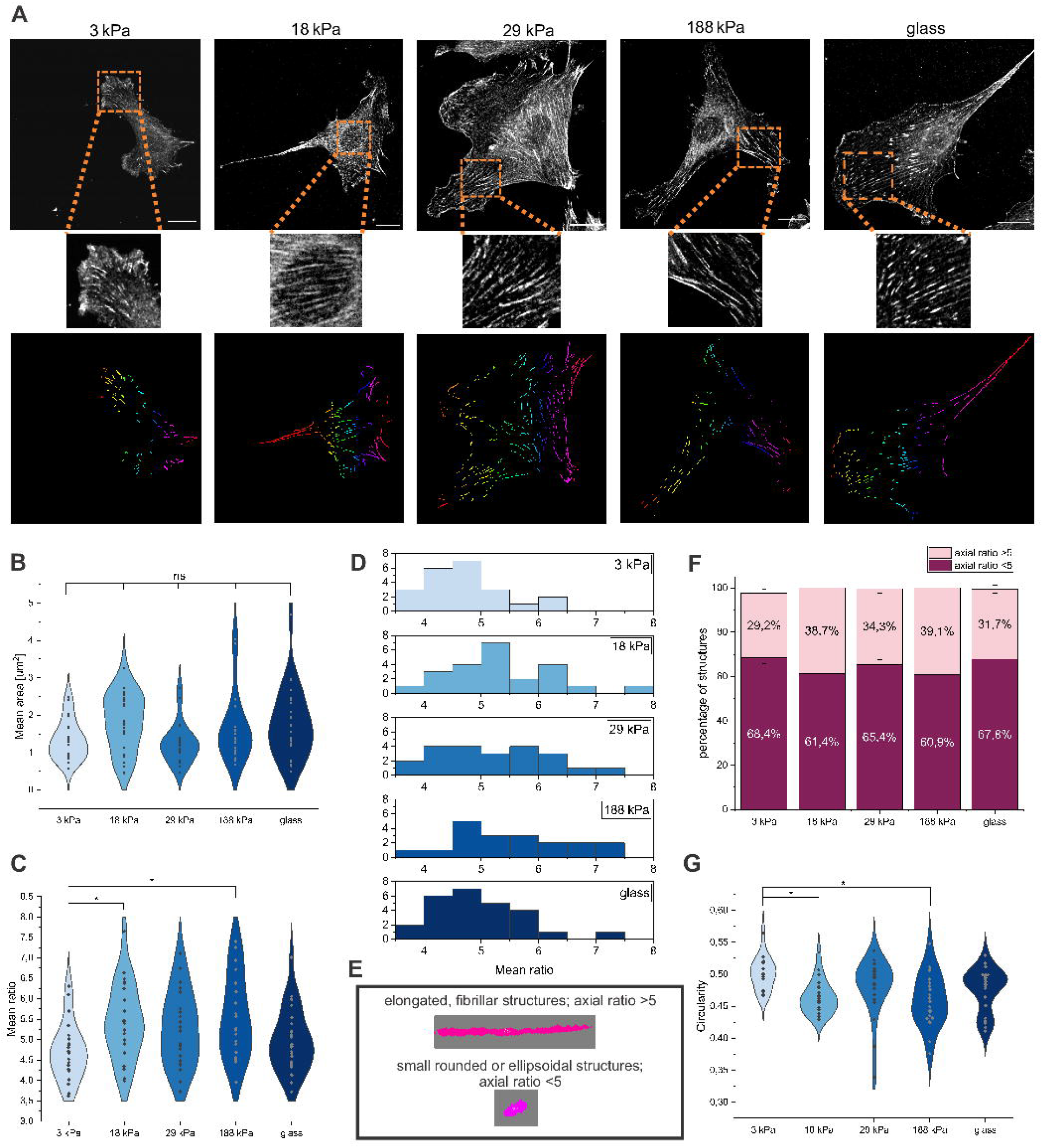
Analysis of talin1 adhesion structures. **A**. Representative images of spatial distribution of talin1 in cells seeded on glass or elastic polyacrylamide gels of different elasticity (3-188 kPa). Top panels show confocal images of talin1 (scale bar: 20 µm). Below (middle panels) show the enlarged interesting fragments of confocal images. The lower panels show images of segmented structures after analyzing software. **B**. Mean area of structures for cells seeded on substrates of different elasticities. **C**. Mean axial ratio of structures for cells seeded on substrates of different elasticities. **D. E. F**. Bar charts comparing percentage of elongated (ratio >5) and rounder (ratio <5) structures in cells seeded on substrates of different elasticity. **G**. Mean circularity of structures for cells seeded on substrates of different elasticities. Statistical analysis: Kruskal-Wallis ANOVA with Dunn’s post-hoc tests, * p ≤ 0.05.

While we showed that the average adhesion structure area remained relatively constant across conditions (ranging 1.3–1.8 µm^2^) (Fig. 3B), the shape varied only slightly depending on the type of substrate. Adhesions on glass resembled “smeared teardrops”, whereas on PA gels looked more like “intermittent fibers”. However, we noticed that our quantitative measurements alone did not fully capture the difference between “eye-drops” and “fibers” shapes of adhesion structures.

Comparing distributions (Fig. 3C) and histograms of mean axial ratio (Fig. 3D) for adhesion structures on different substrates, we observed that at 18 kPa and 188 kPa, they exhibit slightly higher, yet statistically significant, values compared to 3 kPa substrates. We did not significant difference for other conditions. Moreover, structures on 18 kPa, 29 kPa, and 188 kPa gels exhibited a slightly higher proportion of adhesions with an axial ratio >5 (Fig. 3E-F), reaching up to ∼39% on 188 kPa gels, supporting previous observations. Interestingly, the percentage of elongated adhesions on soft 3 kPa substrates (29.2%±1,8%) was similar to glass (31.7%±1,8%). That may suggest that intermediate stiffness promotes adhesion elongation. Circularity analysis showed that adhesions on soft 3 kPa gels were the most rounded (0.492 ± 0.007), while those on stiffer 188 kPa substrates were more elongated (0.459 ± 0.009) (Fig. 3G). We repeated talin1 staining in CRFK feline kidney epithelial cells, receiving similar talin1 organization (Supplementary Fig. 1).

Moreover, talin1 organization in cells on soft ECM, to us, resembled fibronectin, tensin1, and active α5β1 integrin organization in cells, as shown by Barber-Pérez and colleagues (Barber-Pérez et al., 2020). Thus, in further steps of our study, we concentrated on changes in tensin-3 and fibronectin organization.

### ECM stiffness does not influence talin1-tensin3 colocalization

Tensin3 is more specific to fibrillar adhesions, compared to tensin1 and tensin2 (Clark et al., 2010). Moreover, Atherton and colleagues (Atherton et al., 2022) showed recently the importance of the talin1-tensin3 interaction in fibrillogenesis and fibrillar adhesion formation. Thus, we aimed to investigate whether ECM stiffness mediates the colocalization of these proteins and, consequently, their potential interaction. We transiently expressed a tensin3-mEmerald construct in NIH/3T3 cells. After 24 h, we plated them on fibronectin-coated (10 µg/ml) polyacrylamide substrates of 3 kPa, 29 kPa, 188 kPa, and glass. We incubated cells for another 20h, fixed them, stained for talin1, and imaged them using confocal microscopy as described earlier. Using home-written colocalization software, we analyzed colocalization between talin1 and tensin3 (Fig. 4A). We observed intermediate-to-high correlation between talin1 and tensin3 in all our samples, yet we did not find significant differences in Pearson’s coefficient values between samples cultured on polyacrylamide substrates. Interestingly, we observed a difference between glass and all other samples (Fig. 4B).

**Fig. 4.**
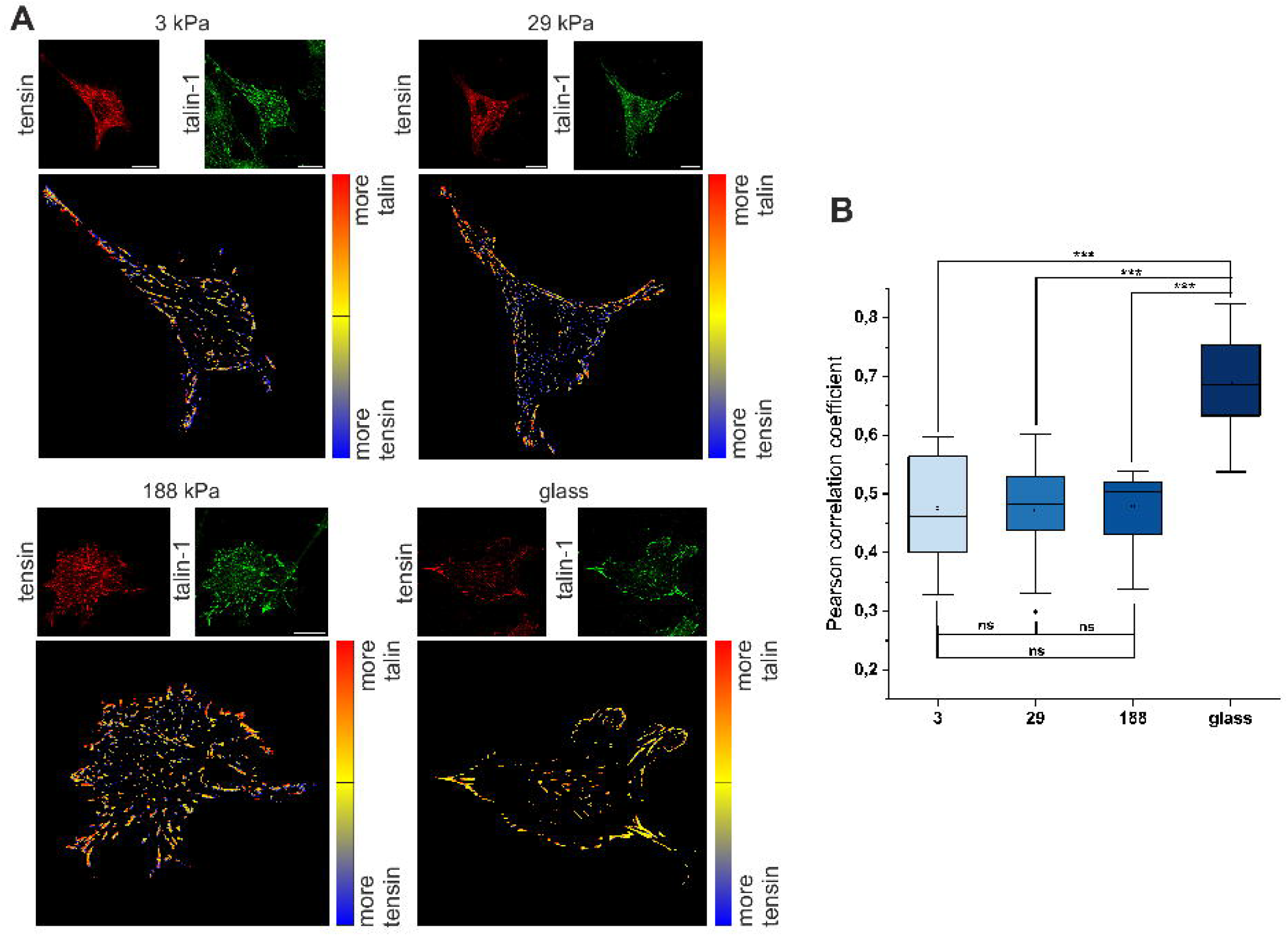
Analysis of talin1-tensin3 colocalization in cells seeded on glass or elastic polyacrylamide gels of different elasticity (3-188 kPa). **A**. Representative images of talin1-tensin3 colocalization in cells. Small panels show distributions of talin1 or tensin3 in cells. Big panels show distributions of the Pearson correlation coefficient-based colocalization within the cell. **B**. Comparison of the mean Pearson correlation coefficient between conditions. Statistical analysis: Kruskal-Wallis ANOVA with Dunn’s post-hoc tests, *** p ≤ 0.001.

### Soft substrate promotes fibronectin secretion in mouse fibroblasts

Finally, we wanted to check how substrate stiffness influences fibronectin organization with respect to talin1. First, we examined whether substrate stiffness influences fibronectin expression in NIH/3T3 mouse embryonic fibroblasts. We plated cells on polyacrylamide substrates of varying stiffnesses embedded with 200 nm fluorescent Dark Red FluoSpheres and coated with 10 µm/ml fibronectin conjugated to HiLite488 to visualize initial coating, and differentiate matrix remodeling from de novo ECM formation. After 24h of culture, we fixed cells with 4% paraformaldehyde in PBS and stained with pan-fibronectin monoclonal antibody and Hoechst 33342 for nucleus visualization. We noticed substantial remodeling of the initial coating on glass compared to practically no remodeling on the PA substrates (Fig. 5A). We attribute this to a different coating procedure, where in the case of glass, fibronectin binds to the substrate via physisorption, while for the PA, it binds covalently via physisorption. Due to high differences in the remodeling, we decided to exclude glass from further analysis. Quantification of the amount of secreted fibronectin under nuclei of cells showed that cells on softer substrates secreted more fibronectin than those on stiffer ones (Fig. 5B).

**Fig. 5.**
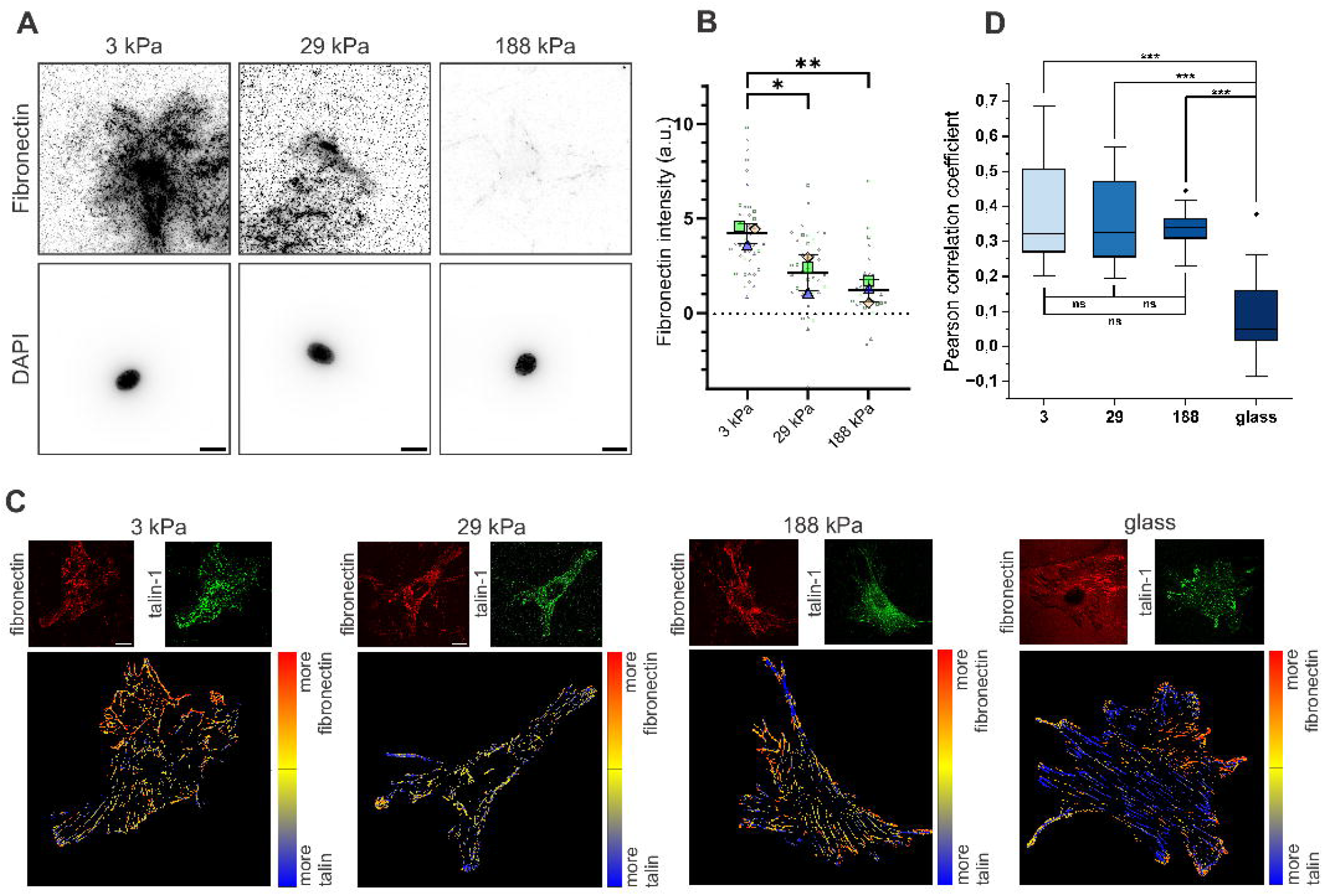
Analysis of fibronectin secretion and talin1-fibronectin colocalization in cells seeded on glass or elastic polyacrylamide gels of different elasticity (3-188 kPa). **A**. Representative images of fibronectin staining for cells plated for 24h on ECM of various stiffnesses, corrected for background and initial coating fluorescence. Additional signal coming from fluorescence beads embedded in the substrate was further accounted for in quantification. **B**. Quantification and comparison of Fibronectin deposition by cells cultured on ECM of different stiffnesses. Statistical analysis: One-way ANOVA with Tukey’s post-hoc tests, *** p ≤ 0.001. **C**. Representative images of talin1-fibronectin colocalization in cells. Small panels show distributions of talin1 or fibronectin in cells. Big panels show distributions of the Pearson correlation coefficient-based colocalization within the cell. **D**. Comparison of the mean Pearson correlation coefficient between conditions. Statistical analysis: Kruskal-Wallis ANOVA with Dunn’s post-hoc tests, *** p ≤ 0.001.

Next, we performed a colocalization experiment between talin1 and fibronectin as we described previously (Fig. 5C). When on glass, we did not see any colocalization; for polyacrylamide substrates, we saw an intermediate strong colocalization independent of substrate stiffness (Fig. 5D). We also investigated colocalization between actin and fibronectin, but we did not see any significance (Supplementary Fig. 2).

## Discussion

Our studies build upon and extend prior works in mechanobiology that linked ECM stiffness to changes in cell behavior and adhesion composition. For example, Oakes and colleagues (Oakes et al., 2018) showed that mouse embryonic fibroblasts on soft substrates switch from a spread, lamellipodial migration mode to a filopodial-driven mode. Similarly, Barber-Pérez and colleagues (Barber-Pérez et al., 2020) demonstrated the mechano-responsiveness of fibrillar adhesion components like tensin1, depending on ECM stiffness. These studies established that a matrix stiffness can reprogram both cell motility and the organization of adhesion structures. However, no study comprehensively examined how ECM stiffness influences talin1 distribution. Our study addresses this gap, revealing new insights into how talin1-mediated adhesion structures adapt to matrix stiffness.

Consistent with previous reports, we observed pronounced morphological changes on soft ECM: cells cultured on intermediate stiffness (around 0.5–11 kPa) displayed reduced spreading, higher edge dynamics, and numerous filopodial protrusions, resembling the behavior noted by Oakes and colleagues, and Pelham and Wang (Oakes et al., 2018; Pelham and Wang, 1997). Beyond these morphology changes, we discovered that talin1 organization on the ECM of different stiffnesses follows different patterns than we anticipated. Similarly to paxillin and vinculin (Han et al., 2021; Oakes et al., 2018; Pelham and Wang, 1997), on stiffer substrates, talin1 was mainly concentrated in peripheral adhesions, whereas on softer substrates talin1 became more evenly distributed throughout the cell (Fig. 1-2). Moreover, quantitative analysis showed a flatter talin1 intensity profile from the cell edge inward on soft ECM, compared to a steep peripheral enrichment on stiff ECMs (Fig. 2). On the other hand, on PA substrates we observed elongated intermittent fiber-like talin1-rich structures, distinct in shape from the ones in cells plated on glass where it resembled a “smeared teardrop” shape. Several previous studies estimated the upper mechanosensitive threshold to be around 20-50 kPa (Barber-Pérez et al., 2020; Sun et al., 2016). By using 188 kPa PA substrates, we expect that, for cells, these substrates will mechanically resemble a glass substrate. Thus, this indicates that there are additional factors influencing talin1-adhesions beyond elasticity. The morphologies of these “fibers” on PA gels were only marginally affected by PA rigidity (Fig. 3), which was of most surprise to us, as earlier studies led to the assumption that changes in talin1 adhesion organization should be similar to those of paxillin or vinculin (Han et al., 2021; Oakes et al., 2018; Pelham and Wang, 1997). The highest discrepancy that we observed comes from the work of Han and colleagues, where they studied talin1 in cells plated on high-refractive-index silicon substrates (Gutierrez et al., 2011). Different substrate types used by Han et al. and us may help explain this discrepancy. Both substrate types were covalently coated with ECM ligands. Thus, we believe the substrate coating was comparable in both cases. However, silicon substrates are purely elastic, whereas PA gels exhibit slight viscoelastic behavior due to their poroelastic nature (Casares et al., 2015). Adebowale and colleagues (Adebowale et al., 2021) showed that ECM viscoelasticity promotes filopodia formation, which might explain the high number of protrusions observed on soft PA gels. Interestingly, talin1 has been shown to regulate filopodia formation through an RIAM-dependent pathway (Lagarrigue et al., 2015; Miihkinen et al., 2021). Soft substrates and the viscoelastic force relaxation are expected to prevent talin1 from unfolding, thereby preserving talin1’s RIAM-binding site (Goult et al., 2021). In turn, this preserved site promotes talin1-RIAM interactions and may lead to subsequent filopodia formation. At the same time, RIAM stabilizes talin1 at the membrane (Lagarrigue et al., 2015). This mechanism may explain the formation of extended talin1 adhesion structures on soft PA substrates.

Notably, the “fiber-like” morphology of talin1 on PA gels resembles that of a talin1^R6–>R13^ mutant, which lacks actin binding at the C-terminal actin-binding site (ABS3) but retains talin1’s ability to dimerize (Driscoll et al., 2020). This suggests lower engagement of ABS3 in force-dependent unfolding of talin1, and preservation of RIAM- and tensin-binding sites at R11 talin1’s rod domain (Atherton et al., 2022; Baster et al., 2025; Goult et al., 2021). Moreover, these elongated adhesions resemble fibrillar adhesions formed on PA hydrogelsgels (Barber-Pérez et al., 2020), suggesting a strong relation to integrin α5β1 and tensin1.

Despite talin1’s and tensin3’s well-known roles as a mechanotransducers, we were surprised by the high level talin1–tensin3 colocalization, and that it appears not to be regulated by ECM stiffness; across substrates from 3 kPa up to 188 kPa, we detected no significant differences in talin1–tensin3 colocalization levels (Fig. 4). This finding suggests that talin1 and tensin3 interaction does not depend on the extracellular rigidity. Atherton and colleagues (Atherton et al., 2022) recently highlighted the importance of the talin1–tensin3 interaction for fibronectin fibril formation, implying that this interaction is biologically significant and likely robust. Our data support that view: talin1–tensin3 coupling seems to be a fundamental aspect of adhesion biology. However, again, we did observe a difference when comparing hydrogel substrates to glass. Cells plated on glass coverslips showed a higher talin1–tensin3 colocalization than those on any polyacrylamide gel, pointing to the influence of factors other than ECM stiffness.

Another finding of our study is that substrate stiffness inversely regulates fibronectin secretion and assembly by fibroblasts. We found that cells on softer substrates secreted significantly more fibronectin into the extracellular matrix than those on stiffer substrates. A similar effect was observed by Atherton and colleagues (Atherton et al., 2022) for U-2 OS cells. Conversely, Carraher and colleagues (Carraher and Schwarzbauer, 2013) showed an inverse dependence for NIH/3T3 than we observed. This discrepancy may come from a slightly different experimental protocol; in their work, they plated cells at a higher density (from our estimation > 50%), while we plated cells to preserve a single-cell environment. They also supplemented full media with additional fibronectin, when we relied on fibronectin present in FBS. In this case, we believe cell-cell contact may be underlying the observed differences, which implies the importance of cell density in fibrillogenesis. Our results suggest a form of mechanoregulatory feedback: when in a soft environment, cells cannot fully engage in integrating catch-bond (Bangasser et al., 2013; Oakes et al., 2018). Fibroblasts may respond by synthesizing more ECM, providing additional adhesion ligands to reinforce their attachment to the substrate. Interestingly, colocalization between talin1 and fibronectin seems to be independent of substrates’ stiffness, suggesting strongly preserved molecular mechanisms connecting fibronectin, tensin3, and talin1 independently from ECM rigidity.

In our studies, we did not look at as broad a range of stiffnesses as Barber-Pérez and colleagues (Barber-Pérez et al., 2020), but we based on our cell motility studies (Movies 1-10) and the work of Oakes and colleagues (Oakes et al., 2018), which both show high differences in cell behavior and, in the case of Oakes et al., in paxillin-adhesion morphology in the chosen stiffness range. In the work of Barber-Pérez et al., they observed that adhesion length starts decreasing below 5 kPa, which we partially observe as a decreased aspect ratio of adhesion structures and an increase in small circularity in cells on 3 kPa substrates. Nonetheless, talin1 adhesions’ morphology itself does not seem to explain the changes that we observe in cell motility. Yet, we believe that our work, together with other works showing the dependence of the distribution of talin1-binding adhesion proteins like vinculin and paxillin on substrate stiffness (Oakes et al., 2018; Pelham and Wang, 1997; Plotnikov et al., 2012; Zhou et al., 2017), supports the role of talin1 as a mechanotransduction hub (Goult et al., 2018). In this case, we show that talin-1-based adhesion morphology does not depend on stiffness, but rather activation and function, and thus downstream signaling, depends on mechanical stimuli.

## Materials and methods

### Cell culture

NIH/3T3 Tet-off embryonic fibroblasts were originally obtained from Clontech (cat. No 630914, Mountain View, CA, USA). NIH/3T3 Tet-off mCherry mouse embryonic fibroblasts were a gift from Dr. Sławomir Lasota (Jagiellonian University, Poland). NIH/3T3 mouse embryonic fibroblasts (CRL-1658), U-2 OS human osteosarcoma (HTB-96), and CRFK feline kidney epithelial (CCL-94) cell lines were obtained from the American Type Culture Collection (ATCC, Manassas, VA, USA).

NIH/3T3 Tet-off mCherry mouse embryonic fibroblasts and NIH/3T3 cells were cultured in Dulbecco’s Modified Eagle Medium with low glucose (DMEM LG; Biowest, Nuaillé, France). U-2 OS and CRFK cells were cultured in McCoy’s 5A medium (ATCC) and Eagle’s minimum essential medium (EMEM; ATCC), respectively. All media were supplemented with 10% fetal bovine serum (FBS; 10270106, Gibco, Thermo Fisher Scientific, Waltham, MA, USA) and 100 units/mL penicillin and 100 μg/mL streptomycin (Biowest). Cells were incubated at 37°C with 5% CO_2_.

For experimental procedures, cells were seeded on 35 mm glass-bottom dishes (thickness #0; CellVis, Mountain View, CA, USA) at the concentration of 2-2.5 * 10^4^ cells and incubated overnight (for approximately 18-24 hours).

### Polyacrylamide (PA) gels preparation

To prepare the gels, 35 mm glass-bottom dishes (CellVis) were pre-treated with a solution of 3-(Trimethylsilyl) methacrylate (Sigma-Aldrich, St. Louis, MO, USA), glacial acid (99.5%) (Chempur, Piekary Śląskie, Poland) and 96% ethanol (1:1:14) for 30 minutes. The treatment was followed by three washes with 96% ethanol.

PA substrates of varying stiffnesses were prepared by mixing different proportions of 40% acrylamide and 2% bis-acrylamide solutions (Bio-rad, Hercules, CA, USA) in PBS (see Table 1). To initiate polymerization, 0.5% ammonium persulphate (Sigma-Aldrich) and 0.05% tetramethylethylenediamine (Bio-rad) were added to each mixture. For fibronectin secretion analysis we added 1:250 of 0.2 μm in diameter Dark Red Carboxylate-Modified FluoSphere Microspheres (F8807, Invitrogen, Thermo Fisher Scientific). A 22 µl aliquot of substrate solution was applied to the center of glass-bottom dish, covered with an 18 mm glass coverslip pre-coated with Sigmacote (SL2, Sigma-Aldrich) and left to polymerize for 1h at room temperature. After polymerization samples were covered with PBS and the coverslip was carefully lifted. Substrates were then stored at 4°C overnight.

**Table 1.** Proportions of 40% acrylamide, 2% bis-acrylamide and PBS used to prepare elastic substrates of different stiffness.

| Published stiffness [kPa] | Measured stiffness [kPa] | 40% acrylamide [$\mu\text{m}$] | 2% bis-acrylamide [$\mu\text{m}$] | PBS [$\mu\text{l}$] | Reference |
| --- | --- | --- | --- | --- | --- |
| 195 | 188 | 250 | 225 | 25 | (Bangasser et al., 2017) |
| 99 | - | 250 | 125 | 125 | (Bangasser et al., 2017) |
| 40 | 29 | 100 | 120 | 280 | (Tse and Engler, 2010) |
| 20 | 18 | 100 | 66 | 334 | (Tse and Engler, 2010) |
| 11 | - | 125 | 25 | 350 | (Tse and Engler, 2010) |
| 2 | 3 | 50 | 25 | 425 | (Tse and Engler, 2010) |
| 1 | - | 62.5 | 7.5 | 430 | (Tse and Engler, 2010) |
| 0.5 | - | 37.5 | 15 | 447.5 | (Tse and Engler, 2010) |
| 0.2 | - | 37.5 | 7.5 | 455 | (Tse and Engler, 2010) |

Next day, substrates were activated with Sulfo-SANPAH (22589, Pierce, ThermFisher) and incubated with 150 µl of 10 µg/ml bovine plasma fibronectin (2385413, Gibco) or 150 µl of 10 µg/ml fibronectin-HiLyte488 (FNR02-A, Cytoskeleton, Inc., Denver, CO, USA) in PBS for 2 hours in room temperature or overnight in 4°C. Substrates were subsequently washed three times with sterile PBS. Glass-bottom dishes were coated with equivalent protein concentrations for the same incubation period as substrates.

### AFM measurement of gel elasticity

AFM measurements were performed with Bruker JPK NanoWizard 3 atomic force microscope (Bruker, Billerica, MA, USA), and the resulting dataset was processed, analyzed, and visualized using JPK Data Processing (Bruker) and OriginPro 2023 software (OriginLab Corporation, Northampton, MA, USA).

The elastic moduli of selected polyacrylamide substrates were evaluated using the nanoindentation method during AFM measurements (Supplementary Fig. 3). Two substrate samples were analyzed for each elasticity. Measurements (force spectroscopy, contact mode) were conducted on at least three separate regions (size of areas: 100×100 µm) of the substrate surface. The maximum setpoint value was 2 nN and approach speed was set to 5 µm/s. A cantilever with a sharp tip (Bruker MLCT-BIO-DC type C, characterized by low thermal sensitivity, with quadratic pyramid tip shape) was used. Contact-based calibration was performed in liquid (distilled water) on a clean microscope slide surface. The elastic modulus (*E*) for each force-distance curve was estimated using Hertz-Sneddon model version appropriate for chosen tip, based on the Bilodeau formula (JPK Instrument AG, 2012). The Poisson’s ratio was set to *ν=0*.*457*, which is a suitable value for polyacrylamide gels (Takigawa et al., 1996).

### Cell immunofluorescence and transient transfection

For talin1 immunofluorescent (co)staining, cells were fixed in ice-cold (−20C) methanol for 3 minutes. After washing in PBS, non-specific binding sites were blocked with 4% BSA (Millipore, Darmstadt, Germany) in PBS for 1 hour. Cells were then incubated overnight at 4°C with primary mouse monoclonal anti-talin-1 antibody (1:100; clone 97H6, Bio-rad) and/or monoclonal anti-fibronectin rabbit antibody (1:200; clone IST-3, Sigma-Aldrich) in 4% BSA in PBS. Then, cells were washed in PBS and incubated with Alexa Fluor 488 goat anti-mouse IgG (H+L), Alexa Fluor 555 goat anti-rabbit IgG (H+L), or Alexa Fluor 647 goat anti-mouse IgG (H+L) antibodies (1:200; Invitrogen) in 4% BSA in PBS for 1 hour.

For actin staining, cells were fixed with 4% paraformaldehyde in PBS for 10 minutes, permeabilized in 0.5% Triton-X 100 in PBS for 5 minutes and incubated with Phalloidin-Atto 488 (1:200; Thermofisher) for 30 minutes. Nuclei were stained with 5µg/ml Hoechst 33342 (H3570; Invitrogen) for 5 minutes.

For transfection, cells were seeded on 35 mm dishes at 70% confluency and transfected with pEGFP tensin3 plasmid (a gift from David Critchley & Kenneth Yamada; Addgene plasmid #105299) (Clark et al., 2010). The transfection was performed using jetOPTIMUS transfection reagent following the manufacturer’s protocol. After overnight incubation, cells were transferred to protein-coated substrates or glass-bottom dishes.

### Confocal microscopy

Cells were imaged using Zeiss LSM 710 confocal scanning microscope, equipped with 40x oil objective (Plan-Apochromat 40×/1.4 NA; Zeiss). Obtained images were denoised using the probabilistic Noise2Void (PN2V) algorithm (Krull et al., 2020).

### Quantitative analysis of focal adhesions properties

In total, 19-23 cells from three samples for each substrate stiffness were analyzed.

A custom application was designed in Matlab to identify adhesion structures in confocal images of fixed samples and calculate various parameters for each object. Parameters included adhesion structures dimensions (area, centroid location, major and minor axis length, perimeter, circularity, orientation) and data regarding their intensity (minimum, maximum and mean intensity, weighted centroid, gradient of intensity). Calculations were performed both for pixel and real size units, based on the image metadata.

User could control the object selection process, manually adjusting regions of interest (ROI) by adding and removing objects. Additionally, objects could be merged or split as necessary.

The software was described in detail elsewhere (Twardawa et. al, *in prep*) and is available online at: https://github.com/patrycja-twardawa/TalinaAppNew.

### Protein colocalization analysis

The correlation between the localization of stained cell structures from two separate image channels (called further red “R” and green “G”) was assessed with a custom script with two primary functionalities: the preparation of colocalization maps and the calculation of the selected quantitative parameter of correlation between channels (Poisson correlation coefficient).

The script allowed for automated image processing (denoising, filtering) and analysis. Visual colocalization maps and Poisson correlation coefficients were calculated for segmented bright objects in the image, separately for the whole cell and the cell nucleus area, defined by the user.

The user could modify the parameters used for processing and segmentation (with a choice of segmentation method), and later adjust the binary maps by excluding unwanted parts from selected ROIs, if there were any.

To analyze protein colocalization across different substrate stiffnesses, images from R and G channels were analyzed for each condition, with two experimental repeats performed for each elasticity. The talin1-tensin3 colocalization included 3 kPa (20 cells), 29 kPa (18 cells), 188 kPa (16 cells), and glass (17 cells). The fibronectin-actin colocalization involved 3 kPa (19 cells), 29 kPa (18 cells), 188 kPa (16 cells), and glass (16 cells). The talin1-fibronectin colocalization analysis comprised 3 kPa (16 cells), 29 kPa (18 cells), 188 kPa (15 cells), and glass (21 cells).

### Statistical analysis

Statistical analysis was performed using OriginPro 2023 and GraphPad Prism 10.

## Acknowledgments

The NIH/3T3 cell line was a gift from Prof. Zbigniew Madeja (Jagiellonian University, Poland), the CRFK cell line was a gift from Prof. Krzystof Pyrć (Malopolska Centre of Biotechnology, Jagiellonian University, Poland). The NIH/3T3 Tet-off mCherry mouse embryonic fibroblasts were a gift from Dr. Sławomir Lasota (Jagiellonian University, Poland). We thank Dr. Martina Lerche (National Heart, Lung, and Blood Institute, National Institutes of Health, USA) for consultation on cellular stainings.

## Author Contributions

Conceptualization: Z.B.; Formal analysis: J.H., P.T., Z.B.; Investigation: J.H., Z.B.; Resources: P.T.; Data curation: J.H., P.T., Z.B.; Writing - original draft: Z.B.; Writing - review & editing: J.H., Z.B.; Visualization: Z.B., J.H.; Supervision: Z.R.; Funding acquisition: J.H, Z.B.

## Data and resource availability

Source data is available from the corresponding authors upon request.

