## Supplementary Materials for "Talin1 adhesions’ morphology is largely unaffected by polyacrylamide substrate stiffness"

**Joanna Hajduk[1,2], Patrycja Twardawa[1,2], Zenon Rajfur[1,3,*], Zbigniew Baster[1,4,*]**

[1] Faculty of Physics, Astronomy and Applied Computer Science, Jagiellonian University, Łojasiewicza 11, 30-348 Cracow, Poland

[2] Doctoral School of Exact and Natural Sciences, Jagiellonian University, Łojasiewicza 11, 30-348 Cracow, Poland

[3]Jagiellonian Center of Biomedical Imaging, Jagiellonian University, Łojasiewicza 11, 30-348 Kraków, Poland

[4]Laboratory for Cell and Tissue Engineering, Department of Biomedical Engineering, Eindhoven University of Technology, 5600 MB Eindhoven, The Netherlands

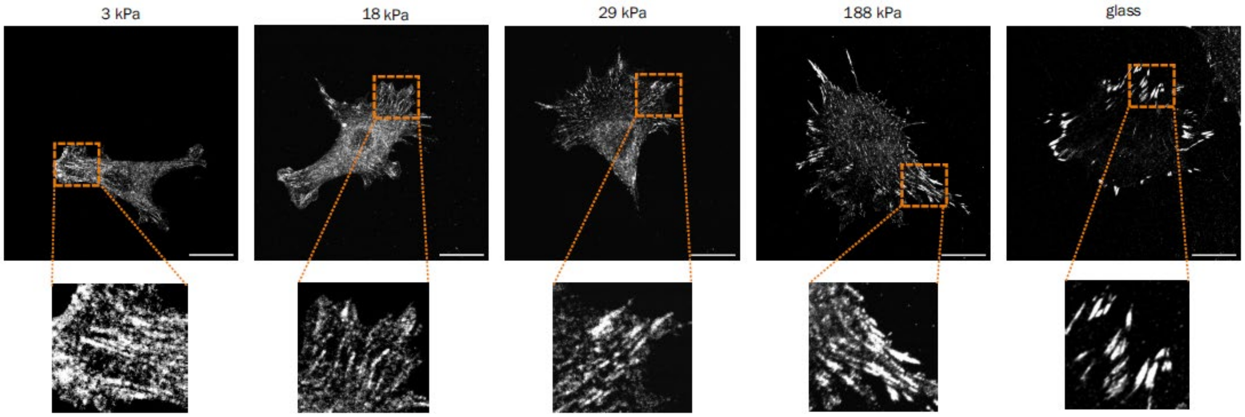


Supplementary Fig. 1. Representative images of the spatial distribution of talin1 in CRF-K cells seeded on glass or elastic polyacrylamide gels of different elasticity (3-188 kPa).

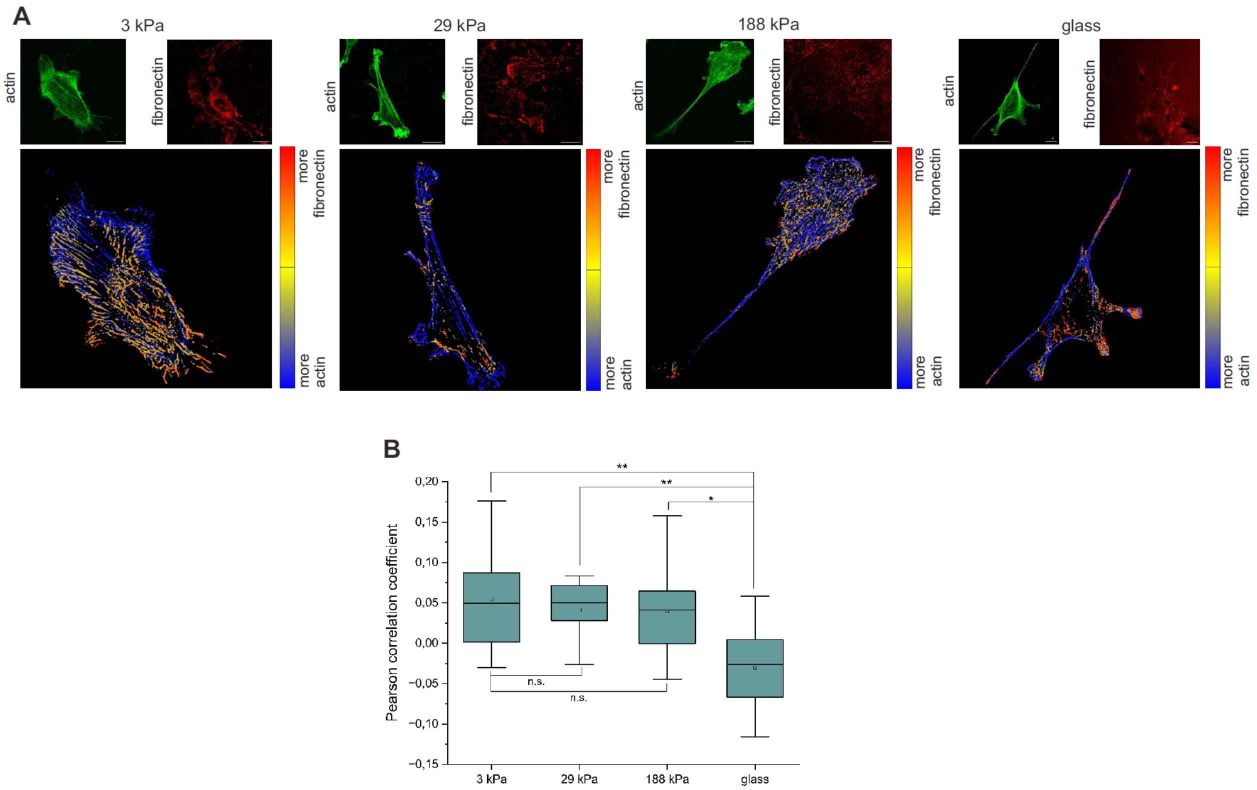


Supplementary Fig. 2. Analysis of actin-fibronectin colocalization in cells seeded on glass or elastic polyacrylamide gels of different elasticity (3-188 kPa). **A.** Representative images of actin-fibronectin colocalization in cells. Small panels show distributions of actin or fibronectin in cells. Big panels show distributions of the Pearson correlation coefficient-based colocalization within the cell. **B.** Comparison of the mean Pearson correlation coefficient between conditions. Statistical analysis: Kruskal-Wallis ANOVA with Dunn's post-hoc tests, * $p \leq 0.05$, ** $p \leq 0.01$.

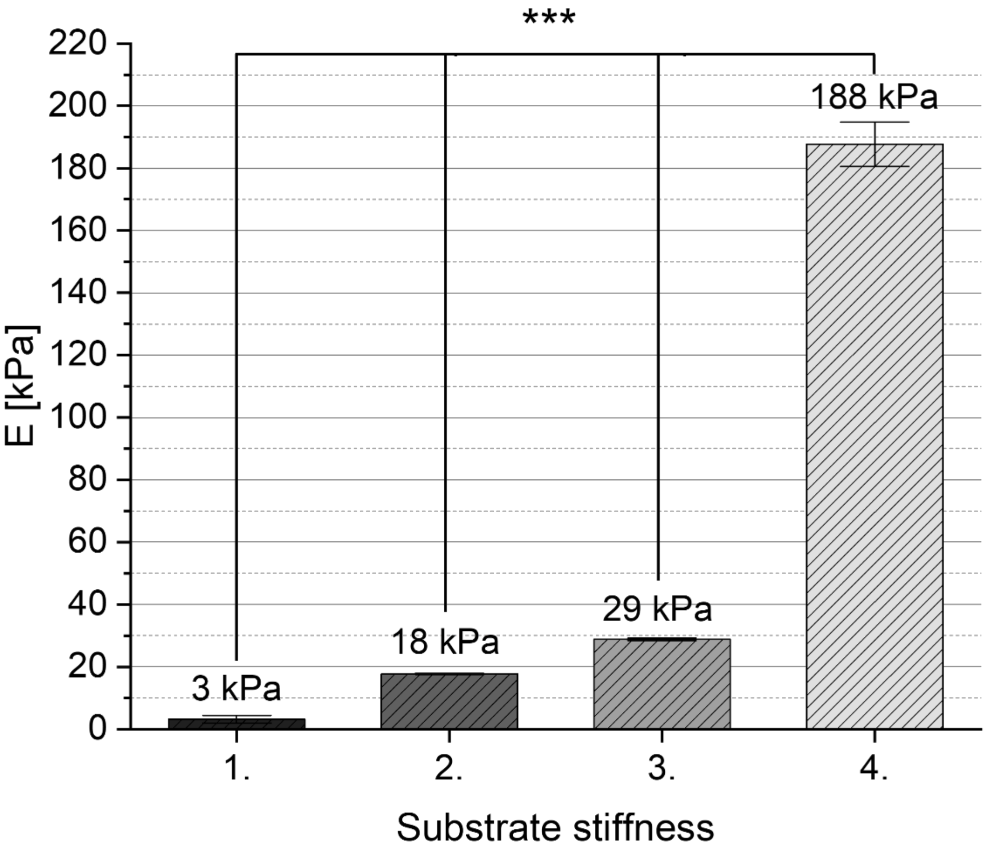


Supplementary Fig. 3. Mean *E* values of polyacrylamide substrates obtained in AFM measurements. Statistical analysis: Kruskal-Wallis ANOVA with Dunn's post-hoc tests (level of statistical significance α ≤0.005). *** $p \leq 0.001$

Movie 1. Mouse embryonic fibroblasts migration on fibronectin-coated glass substrate.

Movie 2. Mouse embryonic fibroblasts migration on fibronectin-coated 188 kPa PA substrate.

Movie 3. Mouse embryonic fibroblasts migration on fibronectin-coated 99 kPa PA substrate.

Movie 4. Mouse embryonic fibroblasts migration on fibronectin-coated 29 kPa PA substrate.

Movie 5. Mouse embryonic fibroblasts migration on fibronectin-coated 18 kPa PA substrate.

Movie 6. Mouse embryonic fibroblasts migration on fibronectin-coated 11 kPa PA substrate.

Movie 7. Mouse embryonic fibroblasts migration on fibronectin-coated 3 kPa PA substrate.

Movie 8. Mouse embryonic fibroblasts migration on fibronectin-coated 1 kPa PA substrate.

Movie 9. Mouse embryonic fibroblasts migration on fibronectin-coated 0.5 kPa PA substrate.

Movie 10. Mouse embryonic fibroblasts migration on fibronectin-coated 0.2 kPa PA substrate.
